# Neural representation of phonological wordform in bilateral posterior temporal cortex

**DOI:** 10.1101/2023.07.19.549751

**Authors:** David O. Sorensen, Enes Avcu, Skyla Lynch, Seppo P. Ahlfors, David W. Gow

**Author notes:** **Corresponding Author:** David W. Gow, Neurodynamics and Neural Decoding Group, Massachusetts General Hospital, 65 Landsdowne Street, rm 219, Cambridge, MA 02139.

## Abstract

While the neural bases of the earliest stages of speech categorization have been widely explored using neural decoding methods, there is still a lack of consensus on questions as basic as how wordforms are represented and in what way this word-level representation influences downstream processing in the brain. Isolating and localizing the neural representations of wordform is challenging because spoken words evoke activation of a variety of representations (e.g., segmental, semantic, articulatory) in addition to form-based representations. We addressed these challenges through a novel integrated neural decoding and effective connectivity design using region of interest (ROI)-based, source reconstructed magnetoencephalography/electroencephalography (MEG/EEG) data collected during a lexical decision task. To localize wordform representations, we trained classifiers on words and nonwords from different phonological neighborhoods and then tested the classifiers’ ability to discriminate between untrained target words that overlapped phonologically with the trained items. Training with either word or nonword neighbors supported decoding in many brain regions during an early analysis window (100-400 ms) reflecting primarily incremental phonological processing. Training with word neighbors, but not nonword neighbors, supported decoding in a bilateral set of temporal lobe ROIs, in a later time window (400-600 ms) reflecting activation related to word recognition. These ROIs included bilateral posterior temporal regions implicated in wordform representation. Effective connectivity analyses among regions within this subset indicated that word-evoked activity influenced the decoding accuracy more than nonword-evoked activity did. Taken together, these results evidence functional representation of wordforms in bilateral temporal lobes isolated from phonemic or semantic representations.

## Introduction

The earliest stages of speech categorization have been solidly grounded by mapping sensitivity to segmental features in the superior temporal gyri through the application of intracranial neurophysiology and neural decoding analyses (Bhaya-Grossman & Chang, 2022; Mesgarani et al., 2014; Oganian & Chang, 2019; Yi et al., 2019). In contrast, it is less clear how this segment-level input is projected to word-level neural representations of any kind (Poeppel & Idsardi, 2022). The goal of the present study is to isolate and localize neural representations of wordform—activity indexing sound patterns that differentiate individual words and mediate the mappings between acoustic-phonetic input and word-specific semantic, syntactic, and articulatory information.

Wordforms play a central role in lexically mediated language-dependent processes ranging from phonetic interpretation, word learning, lexical segmentation, and perceptual learning to sentence processing and rehearsal processes in working memory (Bresnan, 2001; Ganong, 1980; Gathercole et al., 1999; Merriman et al., 1989; Norris et al., 2003). However, questions as basic as how time is represented (Gwilliams et al., 2022; Hannagan et al., 2013), whether wordform representations are episodic or idealized (Pierrehumbert, 2016), morphologically decomposable or holistic (Pelletier, 2012), or fully specified versus underspecified (Lahiri & Marslen-Wilson, 1991) remain topics of vigorous debate. A large body of behavioral and neural research depends on the assumption that nonce or unfamiliar words do not elicit lexical processing because they lack lexical representation of any sort, whereas form representations of words that are in a listener’s lexicon would be stored. Moreover, nonword processing is strongly affected by form similarity to known words (Bailey & Hahn, 2001; Frisch et al., 2000; Gathercole, 1995). Understanding the basis of these similarity effects would clarify the interpretation of results that depend on word-nonword contrasts.

Many studies have examined sensitivity to lexical wordform properties, including cohort size (Gaskell & Marslen-Wilson, 2002; Kocagoncu et al., 2017; Marslen-Wilson & Welsh, 1978; McClelland & Elman, 1986; Zhuang et al., 2014), phonological neighborhood density (Landauer & Streeter, 1973; Luce & Large, 2001; Luce & Pisoni, 1998; Peramunage et al., 2011) and lexical competitor environment (Prabhakaran et al., 2006). These lexical and sublexical effects are considered to play a crucial role in understanding the functional architecture of spoken word recognition and phonotactic constraints that shape wordform representation (Albright, 2009; Hayes & Wilson, 2008; Magnuson et al., 2003; Samuel & Pitt, 2003). An even larger literature has explored BOLD imaging contrasts between words and nonwords (see review by Binder et al., 2009). Several models (Gow, 2012; Hickok & Poeppel, 2007) have synthesized this research, hypothesizing that the bilateral posterior middle temporal gyrus and perhaps the bilateral supramarginal gyrus mediate the mapping between acoustic-phonetic and higher-level representations. While this work coarsely localizes the likely site containing such mediating wordform representations, it does not isolate individual wordforms, which would be an important first step towards characterizing their representation.

Isolating wordforms poses significant challenges. The phonological patterning of wordforms is confounded with the patterning of the phonemes that make up words, making it difficult to discriminate between lexical and segmental representation. This problem is compounded by evidence for the influence of lexical factors on phoneme processing and representation in the brain (Gow et al., 2021; Gow et al., 2008; Gwilliams et al., 2022; Leonard et al., 2016; Myers, 2007). Moreover, auditory words may evoke activation of semantic, articulatory, syntactic, and episodic representations in addition to stored representations of phonological wordform. Finally, evidence that both spoken words and phonotactically legal nonwords briefly activate multiple lexical candidates (Tanenhaus et al., 1995; Zhuang et al., 2014; Zwitserlood, 1989) suggests the need to separate the activation of a target word from that of its coactivated lexical candidates with overlapping phonology.

Neural decoding techniques provide powerful means for investigating the information content of neural signals (Haynes & Rees, 2006; Kriegeskorte & Diedrichsen, 2019; Kriegeskorte & Kievit, 2013). Several studies have used decoding to reconstruct latent information from human brain activity and to understand the relationship between neural representations and cognitive content (Anderson et al., 2019; Choi et al., 2021; Naselaris et al., 2009). The opportunities afforded by these methods have been tempered in part by a tendency to equate decodability with encoding or representation. Decoding analyses reveal the availability of information in neural activity to support a given classification but do not discriminate between latent information and functional representation—which must both capture contrast and influence downstream processing. In addition, functional representation should be localized in a neurally plausible brain region (e.g., an area independently identified as a lexical interface or wordform area). Following Dennett (1987) and Kriegeskorte and Diedrichsen (2019), we suggest that the concept of representation only becomes useful to theory if it can be demonstrated that a representation is plausibly localized and influences downstream processing.

We used a transfer-learning neural decoding and integrated effective connectivity approach, based on ROI-oriented source-reconstructed MEG/EEG activity, to identify and examine wordform representations.

We isolated wordform representations by training classifiers to discriminate between activity evoked by sets of words and nonwords with overlapping phonology, e.g., *pick* and *pid* (neighbors of *pig*) versus *tote* and *tobe* (neighbors of *toad*). We then tested the classifiers’ ability to discriminate between untrained hub words, e.g., *pig* versus *toad*. We reasoned that classifiers trained on activity prior to word recognition or nonword rejection would rely on the segmental overlap between neighbors and hub words, but that classification after word recognition would reflect similarity in global activation patterns associated with the consolidated representation of lexical neighbors. Because neighbors were defined by form similarity rather than semantic similarity, we reasoned that transfer performance would specifically depend on overlap in form representation between hub words and their neighbors. The inclusion of phonologically overlapping nonwords further allowed us to discriminate between lexically mediated classification (which should only occur for words) and sublexical influences (common to both words and nonwords). Finally, to determine whether decoded patterns had causal influences on downstream processing, we used a novel implementation of Granger causality analysis to determine whether within-ROI activation patterns that support decoding in one area influenced decoding performance in downstream processing areas.

## Methods

### Participants

Twenty subjects (14 female) between the ages of 21 to 43 years (mean 29.5, SD = 7.1) participated. All were native speakers of American English and had no auditory, motor, or uncorrected visual impairments that could interfere with the task. Human participation was approved by the Human Subjects Review Board, and all procedures were conducted in compliance with the principles for ethical research established by the Declaration of Helsinki.

### Stimuli

The stimuli consisted of spoken CVC words and nonwords. To limit the potential influence of gross low-level acoustic properties on classification, only stop consonants were used (/b,d,g,p,t,k/). Six words (*pig, toad, cab, bike, dupe, gut*) were chosen as hub words. These were each used to define a set of phonological neighbors. For each hub, we created three word and three nonword neighbors by changing only one phoneme, with the position of the changed phoneme counterbalanced across the three positions. Consonant changes involved either the voice or place feature. Changed vowels were selected to be in close proximity to the hub vowel in F1/F2 space, but this requirement was applied less strictly in order to generate word and nonword tokens. The full set of words and nonwords is included in the Supplementary Material (Table S1). The stimuli were recorded by two speakers, one male and one female. After normalizing the duration to 350 ms and the intensity to 70 dB SPL, a Praat script was used to make additional versions of each token by scaling formant values and mean F0 to create 8 discriminable virtual talkers with different vocal tract sizes (Darwin et al., 2003). This allowed us to both introduce spectral variability within the training set and increase the power of the design.

### Procedure

Simultaneous MEG and EEG data were recorded while the subjects completed a lexical decision task using these stimuli. As the subjects listened to the recorded stimuli, they were asked to determine whether or not the token they heard was a valid English word and signal their judgment via a left-handed keypress on a response pad. The subjects were asked to respond as accurately as possible. Stimuli were presented in 16 blocks, with two non-consecutive blocks assigned to each virtual talker. Within a block, each of the hub words was presented twice and each neighbor once, for a total of 48 trials. Trials were presented in a pseudo-randomized order, with the constraint that the second presentation of a hub word could not directly follow the first presentation. Stimulus presentation was controlled via PsychToolbox for Matlab (Kleiner et al., 2007).

The MEG/EEG data were collected using a Vectorview system (MEGIN, Finland) with 306 MEG channels, a 70-channel EEG cap, and vertical and horizontal electrooculograms. The EEG data were referenced to a nose electrode during the recording. All data were low-pass filtered at 330 Hz and sampled at 1000 Hz. Prior to the testing, the locations of anatomical landmarks (nasion, and left and right preauricular points), four head-position indicator (HPI) coils, the EEG electrodes, and over 100 additional surface points on the scalp were digitized using a FASTRAK 3D digitizer (Polhemus, Colchester, VT). The head position with respect to the MEG sensor array was measured at the start of each block via the HPI coils and was tracked continuously during task performance. In a separate session, T1-weighted structural MRIs were collected from each subject on a 3T Siemens TIM Trio scanner using an MPRAGE sequence.

### Behavioral Analysis

Behavioral accuracy was analyzed using the lme4 (Bates et al., 2012) and lmerTest (Kuznetsova et al., 2017) packages in R (R Core Team, 2023) to perform a logistic mixed-effects analysis of the relationship between accuracy and lexical class (2 levels: Words vs. Nonwords). We ran the full model with word condition as the reference level. Lexical class was treated as a fixed effect. We used random intercepts and slopes for lexical class by participants and random intercepts for lexical class by items. We reported the model estimation of the change in accuracy rate (in log odds) from the reference category for each fixed effect (b), standard error of the estimate (SE), Wald z test statistic (z), and the associated p values.

### MEG/EEG Preprocessing and Source Reconstruction

MEG/EEG data were processed offline using MNE-C (Gramfort et al., 2014) via the Granger Processing Stream software (Gow & Caplan, 2012). Eyeblinks were identified manually, and a set of SSP projectors corresponding to eyeblinks and empty room noise were removed from the MEG data. Epochs from -100 to 1000 ms time-locked to the onset of the auditory stimuli were extracted after lowpass filtering the data at 50 Hz. Epochs with high magnetometer (>100,000 fT) or gradiometer (>300,000 fT/cm) values were rejected. The remaining epochs were averaged across all trials with a correct response for each subject.

Minimum norm estimates (MNE) were calculated for each individual subject to reconstruct event-related electrical activity in the brain (Hämäläinen & Ilmoniemi, 1994). The MEG/EEG source space consisted of ∼10,000 current dipoles located on the cortical surfaces reconstructed from the structural MRIs using Freesurfer (http://surfer.nmr.mgh.harvard.edu/). For the MEG/EEG forward model, a 3-compartment (brain, skull, and scalp) boundary element model was used, with the skull and scalp boundaries obtained from the MRIs. The MRI and MEG/EEG data were co-registered using information from the digitizer and the HPI coils. The MNE inverse operator was constructed with free source orientation for the dipoles. Source estimates were obtained by multiplying the MEG/EEG sensor data with the inverse operator. The source estimates for each individual subject were then brought into a common space obtained by spherical morphing of the MRI data using Freesurfer (Fischl et al., 1999) and averaged to create the group average source reconstruction that was used to perform region of interest (ROI) generation. For the ROI source waveforms used in the decoding and effective connectivity analyses, we calculated noise normalized MNE time courses for each ROI using dynamic statistical parametric mapping (dSPM).

ROIs were generated from the grand average evoked response using the procedures previously described by Gow and Caplan (2012). Briefly, a set of potential centroid locations was generated consisting of the source space dipoles on the cortical surfaces with the highest activation in the time window from 100 to 500 ms after stimulus onset. From those centroids, neighboring dipoles were included into a growing ROI and distant dipoles were excluded from the final set of ROIs based on metrics of similarity, redundancy, and spatial weight. Because our decoding analyses rely on subdividing ROIs, it is beneficial to produce larger ROIs; we thus loosened our redundancy constraints relative to previous studies (Gow & Nied, 2014). This adjustment may increase the rate of type II errors but would not increase the likelihood of false positive results in Granger analyses. This process resulted in a total of 39 ROIs, which were then transformed to each individual subject’s source space.

For decoding analyses, the ROIs were split into 8 approximately equal sized subdivisions. dSPM source time courses were calculated for each subdivision by averaging the source time courses of all the source dipoles within the subdivision. The mean value over the 100-ms pre-stimulus baseline period was subtracted, and the data were vector normalized across subdivisions for each timepoint in each epoch.

### Neural Decoding

Neural decoding analyses with a transfer learning design were conducted to evaluate phonological neighborhood representations using support vector machine (SVM) classifiers (Beach et al., 2021). The SVM classifiers were trained to discriminate neighborhoods using only trials in which the neighbors were presented, then tested on their performance to discriminate the corresponding hub words in a transfer learning design. Pairwise classification was done at each time point within the epoch for all neighborhood pairs.

To increase the robustness of the ROI source waveforms as input to the SVM, bins of 8 trials were randomly selected and averaged within each condition. The random bin assignment was repeated 100 times for each condition. Single timepoints from these bin averages were used as the input to the SVM, and the average classification accuracy across the 100 bin assignments was used as the measure of decoding accuracy for each time point. Performance of these classifiers was then averaged across all subjects and neighborhood pairs. For statistical analysis, the accuracy data were submitted to cluster-based permutation tests. Clusters were defined as consecutive time points of above chance performance (alpha = 0.05; chance performance = 50% for pairwise classification). The observed data were then randomized across time points with 1000 permutations, and the largest cluster within each permutation was taken to form a distribution of cluster sizes. Decoding accuracy was reported as significant if the cluster statistic had p < 0.05.

Our primary decoding analyses contrasted the transfer discrimination of pairs of hub words based on training by exclusively word or nonword neighbors (effect of lexicality). The purpose was to determine the degree to which decoding relied on stored representations of known words as opposed to sublexical overlap present in both neighboring words and nonwords. In addition, to further isolate the effects of sublexical overlap, we compared the decoding of hub word contrasts as a function of positional overlap between the neighbors and hub words. All nonwords in the study partially overlapped with real words, and any decoding based on nonword training could be due to this overlap. Given evidence for the relative importance of onsets in spoken word recognition (e.g., Marslen-Wilson & Tyler, 1980), we hypothesized that overlap between the initial CV-of words and nonwords in the training set and hub words (e.g., the neighbors *pick, pid* and the hub word *pig*) would produce better decoding than overlap involving the final -VC of training words (e.g., *big, tig* and *pig*). To compare between conditions, post-hoc binomial tests were done on two time windows for those ROIs that had at least one significant cluster with time points inside the window in either condition. The early time window (100-400 ms) was chosen to capture incremental phonological correspondences and the late time window (400-600 ms) to capture consolidated lexical effects. The binomial tests compared the total number of time points within the window with above-chance classification accuracy (p<0.05) between the conditions regardless of clustering; results with an FDR-corrected p < 0.05 were considered significant.

### Effective Connectivity Analyses

Effective connectivity analyses follow our previously published Granger causality analysis approach (Gow & Caplan, 2012), with modifications to integrate the results of the decoding analysis as described in Gow et al. (2022). The goal of the modified Granger causality analyses was to identify whether the activation time courses in ROIs that showed decoding could predict (or “Granger cause”) the SVM classifier accuracy time courses in other ROIs. The integration involved substituting the single source waveform activation time course for each ROI (normally used in our Granger analyses) with data relevant to the decoding analyses.

When evaluating the influence of a given ROI on others, the single activation time course for that ROI was substituted by the eight subdivision dSPM time courses used in the decoding analysis. When evaluating how other ROIs influenced a given ROI, the single activation time course for that ROI was substituted by the within-subject neural decoding accuracy time course averaged across all pairwise conditions from a decoding analysis conducted with all neighbors included in the training set. These Granger analyses were limited to those ROIs that showed significantly better transfer decoding for words than nonwords in the late time window (400-600 ms).

Representations, formally defined, require that the activity not only decode but also be related to a functional outcome (Dennett, 1987; Kriegeskorte & Diedrichsen, 2019). Our analysis thus focused on influences between lexicality-sensitive decoders to see how these activations reinforce each other.

Specifically, we examined the ability of the estimated activation time courses in a decoding ROI to predict the decoding accuracy in the other decoding ROIs. We shortened the window of analysis to between 400-500 ms because most ROIs did not show significant decoding after 500 ms and we wanted to ensure effective connectivity was based on above-chance decoding. The strength of Granger causality, as quantified by the number of time points with a significant Granger causality index, was compared between word trials and nonword trials using binomial tests.

## Results

### Behavioral Results

Overall mean behavioral accuracy on the lexical decision was 90% (SD=2.8%), and mean reaction time was 959 ms (SD=148 ms). Accuracy was higher for words (92%; *SD*=4.4%) than for nonwords (86% ; SD=6.3%). This difference was statistically significant (*b* = 1.18, *SE* = 0.46, *z* =2.58, *p* =.01).

### Regions of Interest

A set of 39 ROIs associated with overall task-related activation was identified through our process of clustering cortical source locations associated with activation peaks that share similar temporal activation patterns (Figure 1, Table S2). These ROIs were used for neural decoding and effective connectivity analyses.

**Figure 1.**
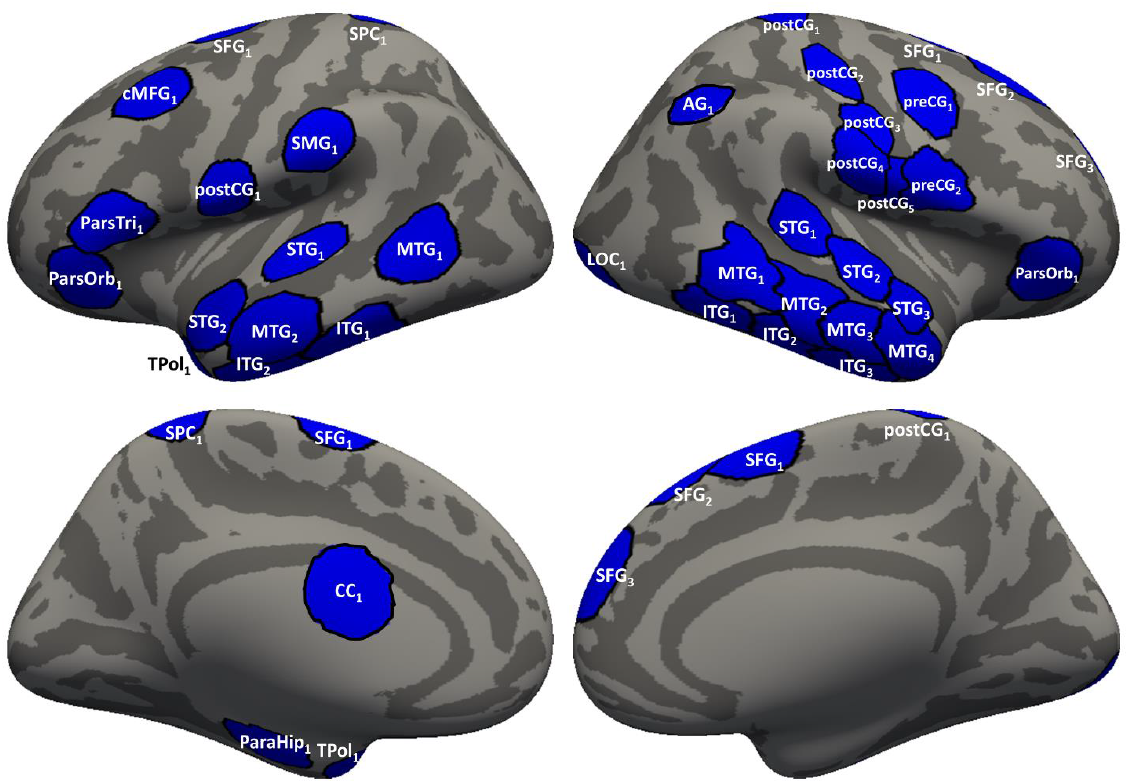
Regions of interest (ROIs) visualized over an inflated averaged cortical surface. Lateral (top) and medial (bottom) views of the left and right hemisphere are shown. For further description of the ROIs, see Table S2.

### Neural Decoding: Lexicality Effects

Figure 2 shows results of the transfer decoding analysis in which SVM classifiers were trained using exclusively either word or nonword neighbors and then tested on their ability to classify the corresponding hub words. Within the entire 1100-ms epoch window, significant clusters of decoding time points were found in 21 of the 39 ROIs. When trained with word neighbors, 20 ROIs produced successful transfer decoding of hub words. When trained with nonword neighbors, 11 ROIs produced successful transfer decoding. Most of the nonword transfer decoding occurred during the early time window when the stimulus could still plausibly be a word. Overall, training with word neighbors resulted in better decoding across more ROIs than training with nonword neighbors.

**Figure 2.**
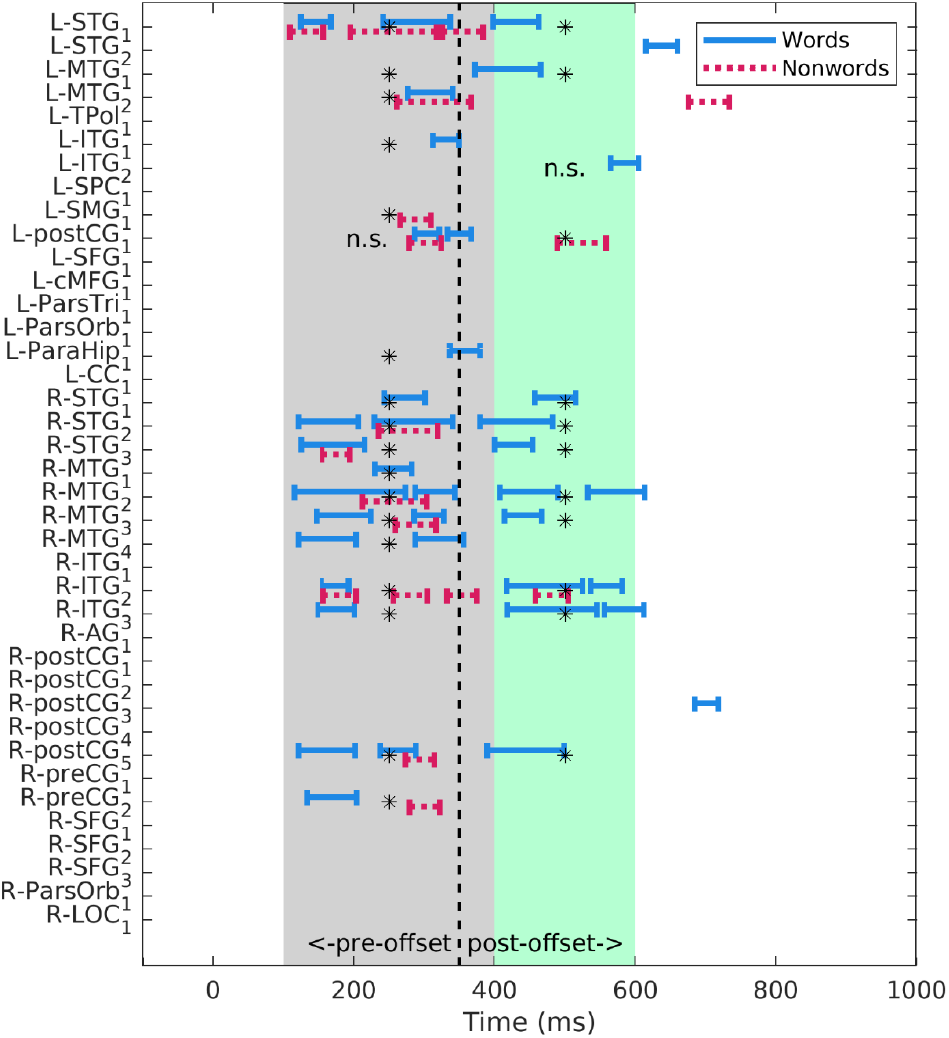
Significant transfer decoding clusters after training with word neighbors or nonword neighbors. This plot summarizes the significant clusters for the words-only (blue) or nonwords-only (red) conditions for the 39 ROIs. The vertical dotted line indicates the offset of the word. Asterisks denote significant differences between the two training conditions (binomial tests, p<0.05) within the early (100-400 ms, grey shading) and the late (400-600 ms, green shading) time windows of interest; n.s.: not significant.

Within the early time window from 100-400 ms, 18 ROIs had significant clusters of decoding time points. Binomial tests indicated better hub word decoding accuracy after word training than nonword training for 13 out of the 18 ROIs (p<0.05, FDR corrected, see Table S3 for detailed stats). Most of these ROIs were located in the bilateral temporal lobe, with the exception of two ROIs in articulatory regions of the central gyrus (R-preCG_2_, R-postCG_5_), and one ROI in the left parahippocampal region (L-ParaHip_1_). Conversely, 4 of the 18 ROIs showed better decoding accuracy after nonword training than word training (L-SMG_1_, L-MTG_2_, L-STG_1_ and R-ITG_2_; p<0.05, FDR corrected).

Within the later window from 400-600 ms, 12 ROIs had significant clusters of decoding time points. 10 of the 12 ROIs showed better decoding accuracy (p<0.05, FDR corrected) in the word than nonword condition, and one ROI (L-postCG_1_) with better decoding accuracy in the nonword than word condition. Regions which produced better decoding accuracy after word training in the late time window included ROIs throughout the right temporal lobe and postcentral gyrus as well as posterior STG and MTG in the left hemisphere. Those 10 ROIs were selected as “decoder ROIs” for the integrated effective connectivity analyses described below.

### Neural Decoding: Positional Effects

We also examined the effects of positional overlap with the hub word regardless of lexicality. Within the entire epoch window, significant clusters of decoding time points were found in 19 of the 39 ROIs (Figure S1). When classifiers were trained by neighbors that shared initial CV-sequences, 14 produced successful transfer decoding of hub words. When trained only by neighbors that shared final -VC sequences, 7 ROIs produced successful transfer decoding of hub words (however, only 3 were within our time windows of interest). For more details, see the Supplementary Materials (Figure S1, Table S4). Overall, better decoding was achieved when training using neighbors with initial than with final overlap.

### Effective Connectivity Analyses

To determine whether local patterns of neural activity influence downstream neural processing, we used a Kalman filter-based implementation of Granger causation analysis that captures directed flow of information between brain regions (Gow & Caplan, 2012; Milde et al., 2010). We focused on interactions among the 10 ROIs that decoded in the late window based on training with word neighbors, contrasting effective connectivity in trials with word versus nonword stimuli. Specifically, we examined whether the event-related neural activity in any one of these decoder ROIs significantly influenced the decoding accuracy in the others within the 400-500 ms time window. We observed that the activity in decoder ROIs influenced the decoding accuracy more in the word than in the nonword condition (Figure 3). For example, significant influences from L-MTG_1_ were all stronger in the word condition than in the nonword condition; and influences to R-MTG_2_ and R-ITG_2_ were stronger in the word condition (R-postCG_5_ being the only exception showing the opposite effect). Most of the other ROIs generally followed these patterns. However, activity in R-STG_2,3_, R-MTG_3_, and R-ITG_2_ influenced the L-STG1 accuracy more in the nonword than in the word condition. Thus, only for L-STG_1_, activity in decoders tended to influence downstream decoding accuracy more in the nonword than in the word condition. These results on the influences between the lexically sensitive decoders indicated that words propagated through the system more than nonwords.

**Figure 3.**
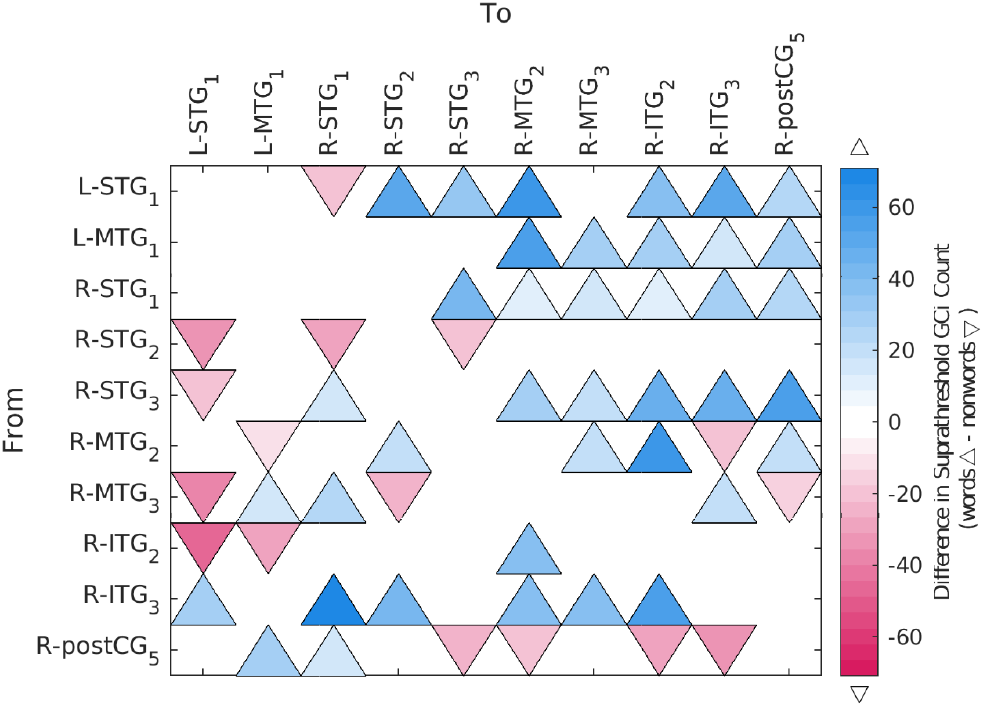
Effective connectivity analysis of transfer decoding ROIs. Matrix representation of Granger causation significantly stronger for words (blue, upward triangles) or nonwords (red, downward triangles) between lexicality sensitive decoding regions. Degree of shading represents the difference in the number of significant Granger causation index timepoints between conditions from 400-500 ms

## Discussion

We undertook this study to isolate wordform representations and localize their neural basis. Word-forms were isolated from segmental overlap in time and from semantic overlap in feature space using phonological neighborhoods. We found significant decoding after training with either word or nonword neighbors during the early window (100-400 ms) consistent with incremental acoustic-phonetic processing before the presentation of the stimulus was complete. In contrast, within the late window of 400-600 ms, decoding occurred mainly after training with word neighbors, suggesting lexical sensitivity in the wordform representation. This late window aligned with the well-characterized event-related potential evidence of lexical processing, including the N400 (Kutas & Federmeier, 2011) and P600 (Kuperberg, 2007). The decoding occurred in a distributed, bilateral network of mostly temporal lobe regions. The effective connectivity analysis showed that the word-evoked activity in these regions preferentially predicted the accuracy of their fellow decoding regions. Our results revealed a bilateral network of temporal lobe areas that were sensitive to both the phonology and lexicality of word-like stimuli and also affected downstream processing.

We achieved these results despite implementing a difficult decoding design. Rather than a leave-n-out cross-validation approach common to many decoding studies (Guggenmos et al., 2018; Hastie et al., 2009), we tested the classifiers in a transfer learning case. The SVMs were trained on one set of words or nonwords and tested on an untrained set of phonologically similar words, based on the assumption that phonologically similar words have overlapping patterns of distributed neural representation. Additionally, our ROI-based approach restricted the signals that the SVM could rely upon to classify epochs. We made these design choices to address qualities necessary to establish a representation. As formally defined by Dennett (1987) and Kriegeskorte & Diedrichsen (2019), for a signal to be a representation it must index the stimulus feature, affect downstream processing, and have a plausible localization. Our integrated approach allowed us to evaluate all three criteria: the decoding analyses tested for feature sensitivity; the effective connectivity analyses examined downstream effects; and the ROI-based approach addressed the location.

Many of the ROIs that decoded hub words based on training with words were in bilateral temporal lobe regions previously implicated a variety of spoken languages functions. Posterior temporal ROIs that decoded in the late window better after training with words than with nonwords included L-MTG_1_, L-ITG_1_, R-ITG_2_, and R-MTG_2_, located in regions that have been implicated in wordform representation. Posterior MTG and adjacent areas have been specifically implicated in wordform representations that mediate the mapping between sound and meaning (Gow, 2012; Hickok & Poeppel, 2007). Imaging studies have shown that activation in these posterior temporal regions is influenced by wordform properties such as word frequency, lexical neighborhood size, lexical enhancement/suppression, phonological similarity, and word-level structural properties (Biran & Friedmann, 2005; Gow et al., 2022; Graves et al., 2007; Prabhakaran et al., 2006; Righi et al., 2009). In addition, damage to posterior temporal regions has been shown to produce deficits in lexico-semantic processing (Axer et al., 2001; Coslett et al., 1987; Goldstein, 1948; Wernicke, 1969).

Anterior temporal ROIs sensitive to wordform included R-ITG_3_ and R-MTG_3,4_, located in regions that have been associated with word retrieval (Abrahams et al., 2003), attention to semantic relations (McDermott et al., 2003), and representing the semantic similarity among concepts (Patterson et al., 2007). Superior temporal ROIs included L-STG_1_ and R-STG_1,2,3_, in regions that have been shown to be sensitive to the processing and representation of the sound structure of language (Hickok & Poeppel, 2007; Mesgarani et al., 2014). Effective connectivity studies of phonotactic repair in phoneme categorization judgments (Gow & Nied, 2014), phonological acceptability judgments (Avcu et al., 2023), and phonotactic frequency effects on lexical decision (Gow & Olson, 2016, 2015; Gow & Segawa, 2009; Gow et al., 2008) have shown the possibility of “referred lexical sensitivity” arising from resonance between acoustic-phonetic representation in superior temporal regions and lexical representation in middle temporal regions and supramarginal gyrus (SMG).

Outside of the temporal lobe, ventral sensorimotor ROIs such as R-postCG_5_ showed decoding for words in both time windows as well as nonwords in the early time window (100-400 ms). Successful decoding in sensorimotor cortex may rely on covert rehearsal. Perception tasks have recruited sensorimotor areas when participants are exposed to foreign phonemes and phoneme segmentation (Callan et al., 2010; Jenson et al., 2014), giving rise to discussion that if a word is unfamiliar (i.e., nonword), it is more likely to be stored in working memory. This argument can also be applied to the fact that the SMG decoded for non-words rather than words (early time window), since the SMG has been shown to play a role in articulatory working memory rehearsal (Gow, 2012; Hickok & Poeppel, 2007). The relationship between the somatosensory cortex and supramarginal gyrus relies on articulatory positions, supported by arguments that somatosensory target and state maps reside in the SMG (Hickok & Poeppel, 2000; Tourville & Guenther, 2011).

The positional analysis (Figure S1; Table S4) showed that word-initial overlap between training and test items supported decoding by more ROIs than did word-final overlap. This aligns with behavioral results that have shown that spoken word recognition relies more heavily on word onsets than offsets (Allopenna et al., 1998; Marslen-Wilson & Tyler, 1980; Marslen-Wilson, 1987). Despite this onset-bias, more ROIs decoded in the word analysis than in the onset analysis, suggesting that overlap at each position contributed to the classification performance in the word condition. This implies that the neural activation patterns reflected parallel activation of stored words with overlapping phonology. Evidence for overlapping neural representation of phonologically similar words suggests a neural basis for lexically mediated “gang effects” supporting a variety of speech and spoken word recognition effects where less activated lexical candidates provide cumulative top-down support based on phonological overlap with input representations. This interpretation is consistent with the claims of the TRACE model (McClelland & Elman, 1986), in which cohort size, length of the target word and phonetic saliency can modulate the intensity of gang effects and their role in onset effects, phonotactic repair, and categorical speech perception (Hannagan et al., 2013).

The results of our integrated effective connectivity analyses strengthen the argument for functional representation of decodable properties. Activity in the decoder ROIs influenced the decoding accuracy in other regions more during word than nonword trials. In particular, ROIs in the posterior MTG (L-MTG_1_ and R-MTG_2_) influenced the accuracy in several other regions more during word than nonword trials. This indicates that the wordform representations propagated information throughout the network. This interpretation is strengthened by the previous work identifying bilateral pMTG as an important wordform area (Gow, 2012; Hickok & Poeppel, 2007). However, we also observed that L-STG1 decoding accuracy was influenced by nonword activity in other decoder ROIs. This may be the result of an intermediate covert step where non-word activity in the right temporal lobe feeds back to left L-STG1 enhancing the representation, repairing it to a word, and producing decoding accuracy. Our reasoning here assumes that this covert feedback/enhancement/repair could be more predictive of decoding accuracy than word activity in other regions, despite the word-evoked activity in L-STG_1_ producing stronger decoding.

Taken together, our results provide evidence of a functional representation of wordform in bilateral temporal lobes. These representations are separated from prelexical phoneme representations in time and independent of semantic relationships. Further work should explore these wordform representations and how they interface to linguistic processes, including accessing semantic content of wordforms.

### Open Practices Statement

The data and materials for this experiment are available by request to pending permanent posting at the Harvard Dataverse (https://dataverse.harvard.edu/). This experiment was not preregistered.

## Acknowledgments

This work was supported by National Institute on Deafness and Other Communication Disorders (NIDCD) grant R01DC015455 (P.I. D.G.). We thank Adriana Schoenhaut and Olivia Newman for their help during data collection. We also thank Dimitrios Pantazis for sharing code used to implement the SVM analyses.

## References

Abrahams, S., Goldstein, L. H., Simmons, A., Brammer, M. J., Williams, S. C., Giampietro, V. P., … Leigh, P. N. (2003). Functional magnetic resonance imaging of verbal fluency and confrontation naming using compressed image acquisition to permit overt responses. Human Brain Mapping, 20(1), 29–40. https://doi.org/10.1002/hbm.10126

Albright, A. (2009). Feature-based generalization as a source of gradient acceptability. Phonology, 26(1), 9–41. https://doi.org/doi:10.1017/S0952675709001705

Allopenna, P. D., Magnuson, J. S., & Tanenhaus, M. K. (1998). Tracking the time course of spoken word recognition using eye movements: Evidence for continuous mapping models. Journal of Memory and Language, 38(4), 419–439. https://doi.org/doi.org/10.1006/jmla.1997.2558

Anderson, A. J., Binder, J. R., Fernandino, L., Humphries, C. J., Conant, L. L., Raizada, R. D., … Lalor, E. C. (2019). An integrated neural decoder of linguistic and experiential meaning. Journal of Neuroscience, 39(45), 8969–8987.

Avcu, E., Newman, O., Ahlfors, S. P., & Gow Jr, D. W. (2023). Neural evidence suggests phonological acceptability judgments reflect similarity, not constraint evaluation. Cognition, 230, 105322.

Axer, H., Keyserlingk, A. G. v., Berks, G., & Keyserlingk, D. G. v. (2001). Supra-and infrasylvian conduction aphasia. Brain and Language, 76(3), 317–331.

Bailey, T. M., & Hahn, U. (2001). Determinants of wordlikeness: Phonotactics or lexical neighborhoods? Journal of Memory and Language, 44(4), 568–591. https://doi.org/10.1006.jmla.2000.2756

Bates, D., Maechler, M., & Bolker, B. (2012). lme4: Linear mixed-effects models using S4 classes. R package version 0.999999-0. In: Vienna.

Beach, S. D., Ozernov-Palchik, O., May, S. C., Centanni, T. M., Gabrieli, J. D., & Pantazis, D. (2021). Neural decoding reveals concurrent phonemic and subphonemic representations of speech across tasks. Neurobiology of Language, 2(2), 254–279.

Bhaya-Grossman, I., & Chang, E. F. (2022). Speech Computations of the Human Superior Temporal Gyrus. Annual Review of Psychology, 73, 79–102.

Binder, J. R., Desai, R. H., Graves, W. W., & Conant, L. L. (2009). Where is the semantic system? A critical review and meta-analysis of 120 functional neuroimaging studies. Cerebral Cortex, 19(12), 2767–2796. https://doi.org/10.1093/cercor/bhp055

Biran, M., & Friedmann, N. (2005). From phonological paraphasias to the structure of the phonological output lexicon. Language and Cognitive Processes, 20(4). https://doi.org/10.1080/01690960400005813

Bresnan, J. (2001). Explaining morphosyntactic competition. In The Handbook of Contemporary Syntactic Theory, M. Baltina and C. Collins (Eds.) (pp. 11–44). Oxford: Blackwell.

Callan, D., Callan, A., Gamez, M., Sato, M.-a., & Kawato, M. (2010). Premotor cortex mediates perceptual performance. Neuroimage, 51(2), 844–858.

Choi, H. S., Marslen-Wilson, W. D., Lyu, B., Randall, B., & Tyler, L. K. (2021). Decoding the real-time neurobiological properties of incremental semantic interpretation. Cerebral Cortex, 31(1), 233–247.

Coslett, H. B., Roeltgen, D. P., Gonzalez Rothi, L., & Heilman, K. M. (1987). Transcortical sensory aphasia: evidence for subtypes. Brain and Language, 32(2), 362–378.

Darwin, C. J., Brungart, D. S., & Simpson, B. D. (2003). Effects of fundamental frequency and vocal-tract length changes on attention to one of two simultaneous talkers. The Journal of the Acoustical Society of America, 114(5), 2913–2922.

Dennett, D. C. (1987). The Intentional Stance. Cambridge MA:The MIT Press.

Fischl, B., Sereno, M. I., Tootell, R. B. H., & Dale, A. M. (1999). High resolution intersubject averaging and a coordinate system for the cortical surface. Human Brain Mapping, 8(4), 272–284. https://doi.org/10.1002/(sici)1097-0193(1999)8:4

Frisch, S., Large, N., & Pisoni, D. (2000). Perception of wordlikeness: Effects of segment probability and length on the processing of nonwords. Journal of Memory and Language, 42(4), 481–496. https://doi.org/10.1006/jmla.1999.2692

Ganong, W. F., 3rd. (1980). Phonetic categorization in auditory word perception. Journal of Experimental Psychology: Human Percepion and Performance, 6(1), 110–125.

Gaskell, M. G., & Marslen-Wilson, W. (2002). Representation and competition in the perception of spoken words. Cognitive Psychology, 45(2), 220–266.

Gathercole, S. E. (1995). Is nonword repetition a test of phonological memory or long-term knowledge? It all depends on the nonwords. Memory & Cognition, 23(1), 83–94.

Gathercole, S. E., Frankish, C. R., Pickering, S. J., & Peaker, S. (1999). Phonotactic influences on short-term memory. Journal of Experimental Psychology: Learning Memory and Cognition, 25(1), 84–95.

Goldstein, K. (1948). Language and language disturbances. New York: Grune & Stratton.

Gow, D., & Nied, A. (2014). Rules from words: Phonotactic biases in speech perception. PloS One, 9(1), 1–12. https://doi.org/10.1371/journal.pone.0086212

Gow, D. W. (2012). The cortical organization of lexical knowledge: A dual lexicon model of spoken language processing. Brain and Language, 121(3), 273–288. https://doi.org/10.1016/j.bandl.2012.03.005

Gow, D. W., & Caplan, D. N. (2012). New levels of language processing complexity and organization revealed by Granger causation. Frontiers in Psychology, 3, 506. https://doi.org/10.3389/fpsyg.2012.00506

Gow, D. W., Jr., & Olson, B. B. (2016). Sentential influences on acoustic-phonetic processing: A Granger causality analysis of multimodal imaging data. Language Cognition and Neuroscience, 31(7), 841–855. https://doi.org/10.1080/23273798.2015.1029498

Gow, D. W., & Olson, B. B. (2015). Lexical mediation of phonotactic frequency effects on spoken word recognition: A Granger causality analysis of MRI-constrained MEG/EEG data. Journal of Memory and Language, 82, 41–55. https://doi.org/10.1016/j.jml.2015.03.004

Gow, D. W., Schoenhaut, A., Avcu, E., & Ahlfors, S. (2021). Behavioral and neurodynamic effects of word learning on phonotactic repair. Frontiers in Psychology, 12, 590155.

Gow, D. W., & Segawa, J. A. (2009). Articulatory mediation of speech perception: a causal analysis of multi-modal imaging data. Cognition, 110(2), 222–236. https://doi.org/10.1016/j.cognition.2008.11.011

Gow, D. W., Segawa, J. A., Ahlfors, S. P., & Lin, F.-H. (2008). Lexical influences on speech perception: A Granger causality analysis of MEG and EEG source estimates. NeuroImage, 43(3), 614–623. https://doi.org/10.1016/j.neuroimage.2008.07.027

Gow, D. W., Avcu, E., Schoenhaut, A., Sorensen, D. O., & Ahlfors, S. P. (2022). Abstract representations in temporal cortex support generative linguistic processing. Language, Cognition and Neuroscience, 1–14.

Gramfort, A., Luessi, M., Larson, E., Engemann, D. A., Strohmeier, D., Brodbeck, C., … Hamalainen, M. S. (2014). MNE software for processing MEG and EEG data. NeuroImage, 86, 446–460. https://doi.org/10.1016/j.neuroimage.2013.10.027

Graves, W. W., Grabowski, T. J., Mehta, S., & Gordon, J. K. (2007). A neural signature of phonological access: distinguishing the effects of word frequency from familiarity and length in overt picture naming [Research Support, N.I.H., Extramural]. Journal of Cognitive Neuroscience, 19(4), 617–631. https://doi.org/10.1162/jocn.2007.19.4.617

Guggenmos, M., Sterzer, P., & Cichy, R. M. (2018). Multivariate pattern analysis for MEG: A comparison of dissimilarity measures. NeuroImage, 173, 434–447.

Gwilliams, L., King, J.-R., Marantz, A., & Poeppel, D. (2022). Neural dynamics of phoneme sequences reveal position-invariant code for content and order. Nature Communications, 13(1), 6606.

Hannagan, T., Magnuson, J. S., & Grainger, J. (2013). Spoken word recognition without a TRACE. Frontiers in Psychology, 4. https://doi.org/10.3389/fpsyg.2013.00563

Hastie, T., Tibshirani, R., & Friedman, J. H. (2009). The elements of statistical learning: data mining, inference, and prediction (Vol. 2). New York: Springer.

Hayes, B., & Wilson, C. (2008). A maximum entropy model of phonotactics and phonotactic learning. Linguistic Inquiry, 39(3), 379–440.

Haynes, J.-D., & Rees, G. (2006). Decoding mental states from brain activity in humans. Nature Reviews Neuroscience, 7(7), 523–534.

Hickok, G., & Poeppel, D. (2000). Towards a functional neuroanatomy of speech perception. Trends in Cognitive Sciences, 4(4), 131–138.

Hickok, G., & Poeppel, D. (2007). The cortical organization of speech processing. Nature Reviews Neuroscience, 8(5), 393–402. https://doi.org/10.1038/nrn2113

Hämäläinen, M. S., & Ilmoniemi, R. J. (1994). Interpreting magnetic fields of the brain: minimum norm estimates. Medical & Biological Engineering & Computing, 32, 35–42.

Jenson, D., Bowers, A. L., Harkrider, A. W., Thornton, D., Cuellar, M., & Saltuklaroglu, T. (2014). Temporal dynamics of sensorimotor integration in speech perception and production: independent component analysis of EEG data. Frontiers in Psychology, 5, 656.

Kleiner, M., Brainard, D., & Pelli, D. (2007). What’s new in Psychtoolbox-3? Perception, 36, 1–16.

Kocagoncu, E., Clarke, A., Devereux, B. J., & Tyler, L. K. (2017). Decoding the cortical dynamics of sound-meaning mapping. Journal of Neuroscience, 37(5), 1312–1319.

Kriegeskorte, N., & Diedrichsen, J. (2019). Peeling the onion of brain representations. Annual Review of Neuroscience, 42, 407–432. https://doi.org/10.1146/annurev-neuro-080317-061906

Kriegeskorte, N., & Kievit, R. A. (2013). Representational geometry: integrating cognition, computation, and the brain. Trends in Cognitive Sciences, 17(8), 401–412.

Kuperberg, G. R. (2007). Neural mechanisms of language comprehension: Challenges to syntax. Brain Research, 1146, 23–49.

Kutas, M., & Federmeier, K. D. (2011). Thirty years and counting: finding meaning in the N400 component of the event-related brain potential (ERP). Annual Review of Psychology, 62, 621–647.

Kuznetsova, A., Brockhoff, P. B., & Christensen, R. H. (2017). lmerTest package: tests in linear mixed effects models. Journal of Statistical Software, 82, 1–26.

Lahiri, A., & Marslen-Wilson, W. (1991). The mental representation of lexical form: A phonological approach to the recognition lexicon. Cognition, 38(3), 245–294.

Landauer, T. K., & Streeter, L. A. (1973). Structural differences between common and rare words: Failure of equivalence assumptions for theories of word recognition. Journal of Verbal Learning and Verbal Behavior, 12(2), 119–131. https://doi.org/doi.org/10.1016/S0022-5371(73)80001-5

Leonard, M. K., Baud, M. O., Sjerps, M. J., & Chang, E. F. (2016). Perceptual restoration of masked speech in human cortex. Nature Communications, 7(13619).

Luce, P. A., & Large, N. (2001). Phonotactics, density, and entropy in spoken word recognition. Language and Cognitive Processes, 16(5), 565–581. https://doi.org/10.1080/01690960143000137

Luce, P. A., & Pisoni, D. B. (1998). Recognizing spoken words: the neighborhood activation model. Ear and Hearing, 19(1), 1–36.

Magnuson, J. S., McMurray, B., Tanenhaus, M. K., & Aslin, R. S. (2003). Lexical effects on compensation for coarticulation: a tale of two systems? Cognitive Science, 27(5), 801–805. https://doi.org/10.1016/s0364-0213(03)00067-3

Marslen-Wilson, W., & Tyler, L. K. (1980). The temporal structure of spoken language understanding. Cognition, 8(1), 1–71.

Marslen-Wilson, W. D. (1987). Functional parallelism in spoken word-recognition. Cognition, 25(1-2), 71–102.

Marslen-Wilson, W. D., & Welsh, A. (1978). Processing interactions and lexical access during word recognition in continuous speech. Cognitive Psychology, 10(1), 29–63.

McClelland, J. L., & Elman, J. L. (1986). The TRACE model of speech perception. Cognitive Psychology, 18(1), 1–86.

McDermott, K. B., Petersen, S. E., Watson, J. M., & Ojemann, J. G. (2003). A procedure for identifying regions preferentially activated by attention to semantic and phonological relations using functional magnetic resonance imaging. Neuropsychologia, 41(3), 293–303. https://doi.org/10.1016/s0028-3932(02)00162-8

Merriman, W. E., Bowman, L. L., & MacWhinney, B. (1989). The mutual exclusivity bias in children’s word learning. Monographs of the Society for Research in Child Development, i–129.

Mesgarani, N., Cheung, C., Johnson, K., & Chang, E. F. (2014). Phonetic feature encoding in human superior temporal gyrus. Science, 343(6174), 1006–1010. https://doi.org/10.1126/science.1245994

Milde, T., Leistritz, L., Astolfi, L., Miltner, W. H., Weiss, T., Babiloni, F., & Witte, H. (2010). A new Kalman filter approach for the estimation of high-dimensional time-variant multivariate AR models and its application in analysis of laser-evoked brain potential. NeuroImage, 50(3), 960–969. https://doi.org/10.1016/j.neuroimage.2009.12.110

Myers, E. B. (2007). Dissociable effects of phonetic competition and category typicality in a phonetic categorization task: an fMRI investigation. Neuropsychologia, 45(7), 1463–1473. https://doi.org/10.1016/j.neuropsychologia.2006.11.005

Naselaris, T., Prenger, R. J., Kay, K. N., Oliver, M., & Gallant, J. L. (2009). Bayesian reconstruction of natural images from human brain activity. Neuron, 63(6), 902–915.

Norris, D., McQueen, J. M., & Cutler, A. (2003). Perceptual learning in speech. Cognitive Psychology, 47(2), 204–238.

Oganian, Y., & Chang, E. F. (2019). A speech envelope landmark for syllable encoding in human superior temporal gyrus. Science Advances(eaay6279).

Patterson, K., Nestor, P. J., & Rogers, T. T. (2007). Where do you know what you know? The representation of semantic knowledge in the human brain [Review]. Nature Reviews. Neuroscience, 8(12), 976–987. https://doi.org/10.1038/nrn2277

Pelletier, F. J. (2012). Holism and compositionality. In M. Werning, W. Hinzen, & E. Maachery (Eds.), The Oxford Handbook of Compositionality (pp.149–174). Oxford: Oxford University Press.

Peramunage, D., Blumstein, S. E., Myers, E. B., Goldrick, M., & Baese-Berk, M. (2011). Phonological neighborhood effects in spoken word production: an fMRI study. Journal of Cognitive Neuroscience, 23(3), 593–603. https://doi.org/10.1162/jocn.2010.21489

Pierrehumbert, J. B. (2016). Phonological representation: Beyond abstract versus episodic. Annual Review of Linguistics, 2, 33–52.

Poeppel, D., & Idsardi, W. (2022). We don’t know how the brain stores anything, let alone words. Trends in Cognitive Sciences.

Prabhakaran, R., Blumstein, S. E., Myers, E. B., Hutchison, E., & Britton, B. (2006). An event-related fMRI investigation of phonological–lexical competition. Neuropsychologia, 44(12), 2209–2221. https://doi.org/10.1016/j.neuropsychologia.2006.05.025

Righi, G., Blumstein, S. E., Mertus, J., & Worden, M. S. (2009). Neural systems underlying lexical competition: An eye tracking and fMRI study. Journal of Cognitive Neuroscience, 22(2), 213–224.

Samuel, A. G., & Pitt, M. A. (2003). Lexical activation (and other factors) can mediate compensation for coarticulation. Journal of Memory and Language, 48(2), 416–434.

Tanenhaus, M. K., Spivey-Knowlton, M. J., Eberhard, K. M., & Sedivy, J. C. (1995). Integration of visual and linguistic information in spoken language comprehension. Science, 268(5217), 1632–1634.

Tourville, J. A., & Guenther, F. H. (2011). The DIVA model: A neural theory of speech acquisition and production. Language and Cognitive Processes, 26(7), 952–981.

Wernicke, C. (1969). The symptom complex of aphasia: A psychological study on an anatomical basis. Proceedings of the Boston Colloquium for the Philosophy of Science 1966/1968,

Yi, H. G., Leonard, M. K., & Chang, E. F. (2019). The encoding of speech sounds in the superior temporal gyrus. Neuron, 102(6), 1096–1110.

Zhuang, J., Tyler, L. K., Randall, B., Stamatakis, E. A., & Marslen-Wilson, W. D. (2014). Optimally efficient neural systems for processing spoken language. Cerebral Cortex, 24(4), 908–918.

Zwitserlood, P. (1989). The locus of the effects of sentential-semantic context in spoken-word processing. Cognition, 32(1), 25–64.

